# Integrative analysis and refined design of CRISPR knockout screens

**DOI:** 10.1101/106534

**Authors:** Chen-Hao Chen, Wei Li, Tengfei Xiao, Han Xu, Peng Jiang, Clifford A. Meyer, Myles Brown, X. Shirley Liu

## Abstract

Genome-wide CRISPR-Cas9 screen has been widely used to interrogate gene functions. However, the analysis remains challenging and rules to design better libraries beg further refinement. Here we present MAGeCK-NEST, which integrates protein-protein interaction (PPI), improves the inference accuracy when fewer guide-RNAs (sgRNAs) are available, and assesses screen qualities using information on PPI. MAGeCK-NEST also adopts a maximum-likelihood approach to remove sgRNA outliers, which are characterized with higher G-nucleotide counts, especially in regions distal from the PAM motif. Using MAGeCK-NEST, we found that choosing non-targeting sgRNAs as negative controls lead to strong bias, which can be mitigated by sgRNAs targeting the “safe harbor” regions. Custom-designed screens confirmed our findings, and further revealed that 19nt sgRNAs consistently gave the best signal-to-noise separation. Collectively, our method enabled robust calling of CRISPR screen hits and motivated the design of an improved genome-wide CRISPR screen library.

## Introduction

The clustered, regularly interspaced, short palindromic repeat (CRISPR)-Cas9 system is a new genome editing technology that becomes prominent in many biomedical research areas. In this system, single guide RNAs (sgRNAs) direct Cas9 nucleases to induce double-strand breaks at targeted genomic regions^1–3^. Based on this system, CRISPR-Cas9 loss-of-function screens can interrogate the functions of coding genes^4–7^ and non-coding elements^8–10^, and generate hypotheses on cell dependency, drug response, and gene regulation in a high-throughput and unbiased manner^11–14^. From a computational biology perspective, several algorithms have been developed to characterize sgRNAs with high specificity and efficiency^15–17^ that can be used in designing CRISPR screen libraries. Algorithms to analyze screening data using either ranking or likelihood approach have also been developed, such as RIGER^18^, RSA^19^, HitSelect^20^, ScreenBeam^21^, casTLE^22^, as well as the MAGeCK/MAGeCK-VISPR/NEST algorithms we previously published^23, 24^. Despite these efforts, methods for designing CRISPR screens and identifying hits from the screens are still being refined from different aspects. First, the number of sgRNAs in the library influences the sensitivity of the screens. Libraries with fewer sgRNAs per gene^4, 16^ often detect fewer statistically significant genes^23^, so algorithms to increase the analysis power of the screens are needed. We have demonstrated in a recent method NEST^25^ that existing biological knowledge such as protein-protein interaction networks can improve CRISPR screen analysis. However, NEST did not directly incorporate such information in the statistical model to improve hit calling from the screens. Second, sgRNAs outliers, or sgRNAs with discrepant behaviors from other sgRNAs targeting the same gene, skew the hit calling results, especially when fewer sgRNAs target each gene. However, rules to predict, detect, and remove outlier sgRNAs in CRISPR screens, and designing sgRNA libraries with high efficiency and specificity, are still lacking.

Here we present an analytical framework MAGeCK-NEST (Fig. 1) which extends our previous MAGeCK-VISPR and NEST algorithms^24,25^. MAGeCK-NEST is able to utilize protein-protein interaction (PPI) information to improve the accuracy and statistical power of hit calling from CRISPR screens with limited number of sgRNAs per gene. Applying MAGeCK-NEST to published screens^5, 11^, we identified and removed outlier sgRNAs and uncovered their sequence features to inform future library design. We also found a strong bias on CRISPR screen gene selection when normalizing read counts with commonly used non-targeting sgRNAs, and proposed an alternative normalization to mitigate such bias. We performed custom-designed screens to validate these findings, and further explored sgRNA design rules that can improve the screening results, including the optimal spacer length for higher cutting efficiencies and better signal-to-noise ratios. Finally, we designed a genome-wide CRISPR/Cas9 screening library based on these new design rules, and demonstrated its performance in identifying known essential genes in different cell types.

**Figure 1.**
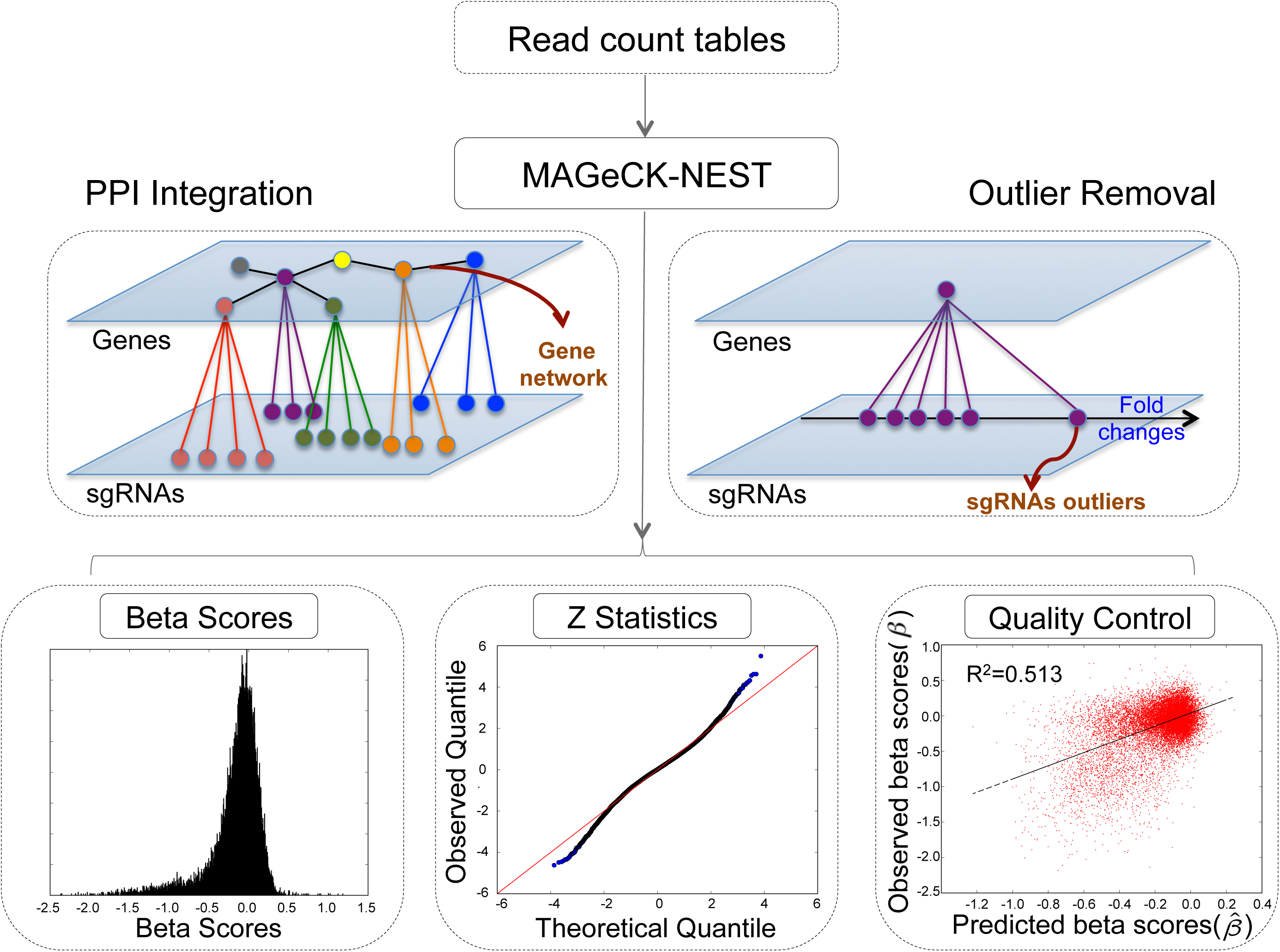
An overview of the MAGeCK-NEST workflow. The input of MAGeCK-NEST is a read count table to record the counts of every sgRNA in all samples. MAGeCK-NEST then builds a hierarchical model based on information from protein-protein interaction (PPI), and removes outliers that have aberrant fold changes compared with other sgRNAs within the same gene. MAGeCK-NEST outputs beta scores (measuring the degree of selections of all genes), p-values, and quality control metrics.

## Results

### The MAGeCK-NEST algorithm

Our laboratory has previously developed algorithms MAGeCK and MAGeCK-VISPR for identifying CRISPR screen hits in different scenarios^23, 24^. In two-condition comparisons, MAGeCK uses a negative binomial model to assess the degree of selections of individual sgRNAs, and adopts robust rank aggregation (RRA) algorithm^27^ to aggregate multiple sgRNAs on a gene to evaluate gene selection. MAGeCK-VISPR^24^ further quantitatively estimates gene selections by optimizing a joint likelihood function of observing the read counts of different sgRNAs with varying behaviors in multiple conditions. The output of MAGeCK-VISPR is a “beta score” for each gene, analogous to the “log fold change” in differential gene expression analysis. We also developed an algorithm, NEST^25^ (Network Essentiality Scoring Tool), to investigate the relationship between gene essentiality and gene network information. We found that the expressions of protein interacting neighbors from the STRING database^20^ can help predict gene essentialities, implicating the potential benefit of further integrating PPI information to improve hit calling from CRISPR screens.

In this study, we propose MAGeCK-NEST that adopts a Bayesian framework to integrate the salient features from MAGeCK-VISPR and NEST (Supplementary Fig. 1). First, MAGeCK-NEST uses MAGeCK-VISPR to estimate the beta score for each gene in a condition. Then for each gene of interest *g*, MAGeCK-NEST estimates an informative prior on the beta score of *g*, based on the weighted average of beta scores on *g*′s PPI neighbors (Fig. 2a). Indeed, there is a good correlation between observed beta score and predicted prior in published screen datasets^11^, with correlation coefficients as high as 0.5, indicating the consistent gene knock-out effects between interacting proteins (Fig. 2b). Finally, MAGeCK-NEST uses the actual CRISPR selection observed on *g* to iteratively optimize the posterior probability likelihood of observing read counts of sgRNAs targeting *g*. A detailed description of MAGeCK-NEST can be found in Methods.

**Figure 2.**
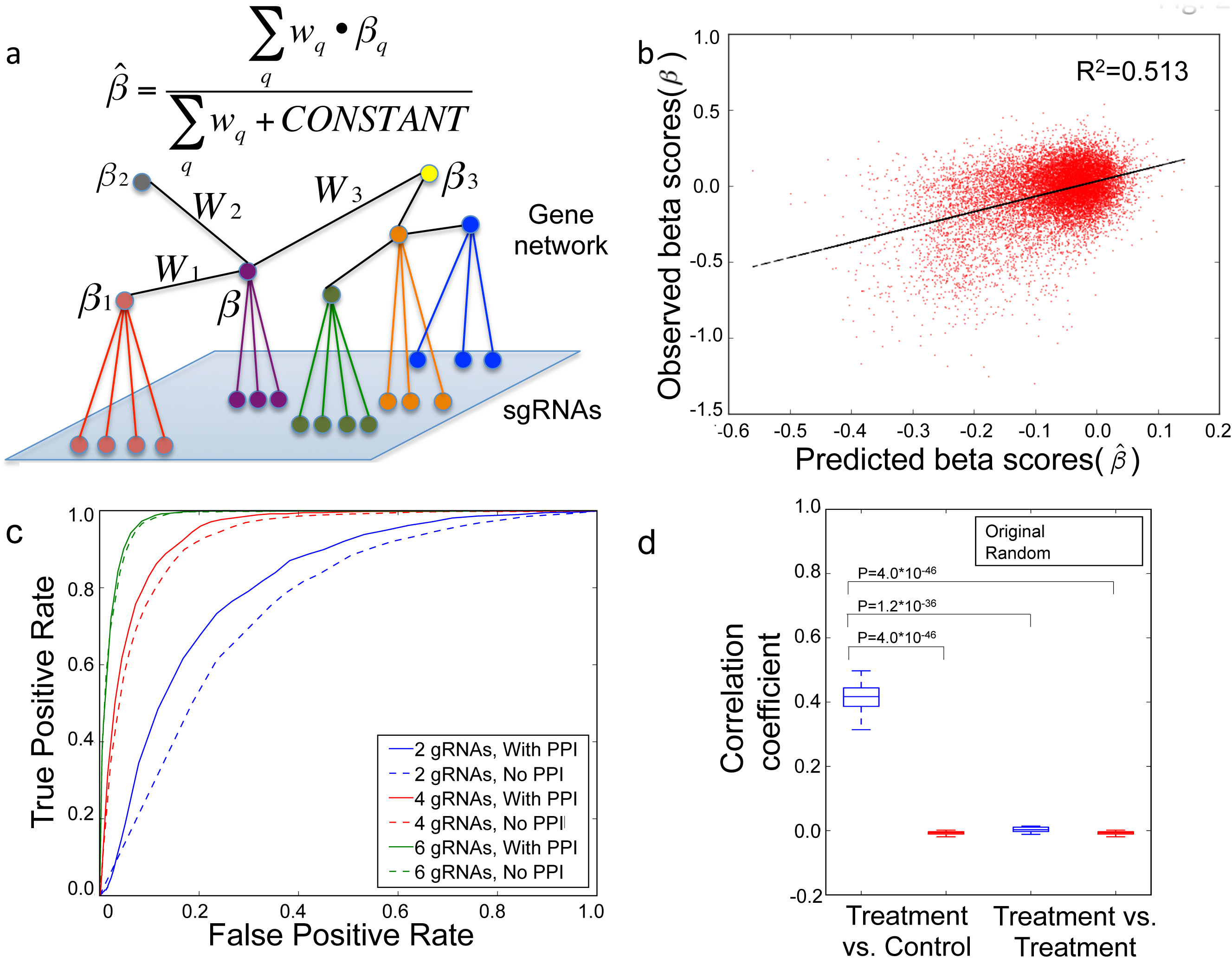
Integrating PPI to call significant genes from genome-wide CRISPR-Cas9 screens and serve as quality control metric. (a) Predicting gene knockout effect using the weighted average of interacting genes in PPI. (b) The correlation between the PPI-predicted and observed beta scores. (c) The receiver operating characteristic (ROC) curves for identifying significant genes with or without PPI using different numbers of sgRNAs. The “gold standard” genes are defined as those that are statistically significant when 10 sgRNAs are used. (d) The distribution of correlation coefficients between predicted and observed beta scores, from 33 “effective screens” (4 treatments vs. 1 corresponding control condition) and 29 “ineffective screens” (2 treatments vs. 2 treatments). The interacting genes in PPI (blue) as well as randomized PPI (red) are used for calculating the correlation. P values were calculated using two-sided Student’s t-test.

The major advantage of the Bayesian framework is that the prior is negligible when the behaviors of sgRNAs on a gene are consistent, but it could potentially play a critical role when the behaviors from sgRNAs are inconsistent^28^. Also, incorporating the informative prior from PPI would not sacrifice specificity (Supplementary Fig. 2). To evaluate MAGeCK-NEST, we down-sampled sgRNAs from a CRISPR screen dataset containing 10 sgRNAs per gene^11^, and compared the number of significant genes called with or without PPI. Using genes called with 10 sgRNAs as gold standard, we found integrating PPI could give better predictive power even with fewer available sgRNAs, as indicated by the higher Area Under Curve score in Receiver Operating Characteristic (Fig. 2c). One such example is the gene PRSA, an mitochondrial gene that is identified as essential when 10 sgRNAs are used^11^ by MAGeCK/VISPR but would be missed with 4 sgRNAs, and this essential gene could be rescued by applying the prior (Supplementary Fig. 3).

**Figure 3.**
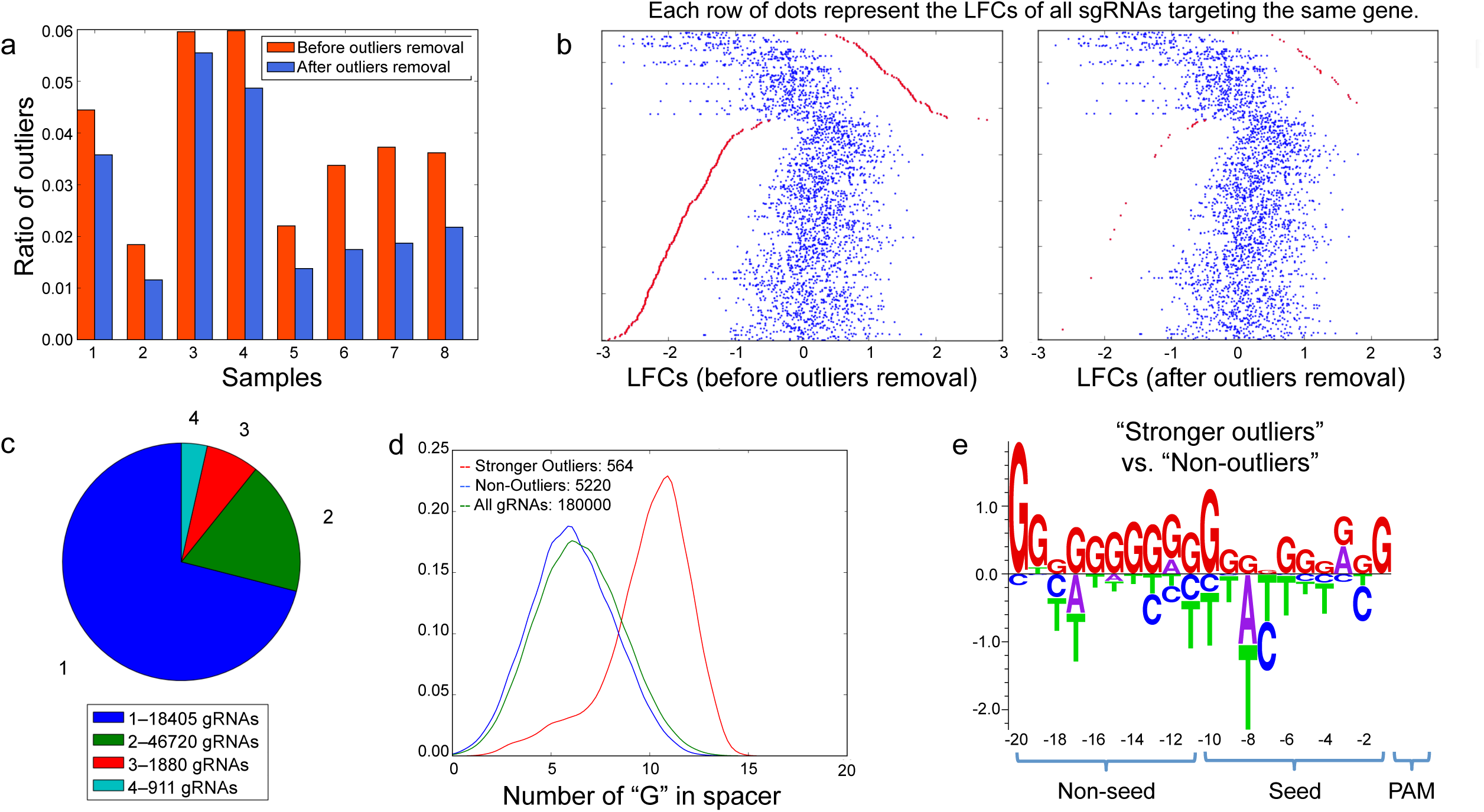
Identifying and characterizing sgRNAs outliers using MAGeCK-NEST. (a) The ratio of sgRNAs that fall beyond 2 standard deviations from the mean before and after outlier removal in published screening data^4, 5, 11^. (b) Identifying and removing aberrantly stronger outliers (red dots). Each row of dots represents the log fold changes (LFCs) of sgRNAs targeting the same gene. (c) The numbers of sgRNAs that are repeatedly identified as outliers in four screening cell lines in a public screening dataset^11^. (d) The G-nucleotide counts of sgRNAs in three groups: stronger outliers (red), non-outliers (blue), and all sgRNAs (green). (e) The sequence features of stronger outliers versus non-outliers derived by elastic-net regression. The “seed” and “non-seed” regions are defined as a 10-nucleotide window proximal to and distal from the PAM motif, respectively. The data of Fig. 4b-e come from a public screening dataset^11^.

Quality control (QC) is critical to ensure that data from CRISPR screens is of high quality and could be evaluated at different levels^23^. Since interacting genes often show similar selection in a screen (Fig. 2b), this information can be used as a quality control (QC) metric in genome-wide screens. To test whether the correlation between observed and predicted beta scores (derived from interacting genes using PPI) can reflect screen quality, we calculated the correlation coefficients of these gene pairs in different settings. These include 33 “effective” screen comparisons (*i.e.,* treatment vs. corresponding control conditions, where these pairs should have positive correlation) and 29 “ineffective” screen comparisons (*i.e*., replicated conditions where no genes should be selected)^29^. In each setting, we also compared the results using the original STRING PPI and randomized PPI as a control. A higher distribution of correlation coefficients in PPI gene pairs is observed in “effective” screens with the original PPI, compared to other three groups (Fig 2d). This suggests that the knockout effects show agreements between interacting genes, which could be used to evaluate screen quality.

### sgRNAs outlier identification, removal, and characterization

Different sgRNAs targeting the same gene may result in varying phenotypes or selection levels in the screen due to different cleavage and repair efficiencies, local chromatin structure, protein domains, and potential off-target effects, *etc*^31–33^. Some sgRNAs with outlier phenotypes compared with other sgRNAs on the same gene, regardless of the causes, may yield false positive or false negative calls in the screens (e.g., the outlier presented in Supplementary Fig. 3, Supplementary Fig. 4). From published screens ^4, 5, 11^, 2-8% of sgRNAs has log fold change (LFC) over 2 standard deviation (STDEV) away from LFC of other sgRNAs targeting the same gene (Fig. 3a), suggesting that their effect is not ignorable. Some outliers behave consistently in multiple screen conditions^11^ (Supplementary Fig 4), suggesting that the discrepant phenotypes could arise from intrinsic features of the sgRNA in addition to random variances in the experiments.

In MAGeCK-NEST, we implemented an approach to identify such outliers, which tests whether one sgRNA has big effect on the beta score estimates of a gene or the likelihood of observing the sgRNA conditioned on the beta score of the gene is low (see Methods). This outlier detection and removal approach can significantly reduce the number of sgRNAs with aberrant LFC on a gene (Fig. 3a-b). In published screens on four leukemia cell lines^11^, nine thousand out of 182K sgRNAs on average were identified as outliers, among which 911 are outliers in all four screens (Fig. 3c). Among these, some outliers have much stronger absolute LFCs compared with other sgRNAs targeting the same gene (Fig 3b, Supplementary Table 1). When examining the sequence features of these strong outliers, we found that they have higher G-nucleotide counts (but lower C-nucleotide counts) that spread across the spacers (Figure 3d, Supplementary Fig. 5). Using elastic net regression^34^ to identify sequence features distinguishing between outliers and non-outliers, we found that outliers contain more G-nucleotides in the 10-nucleotide non-seed region distal from the PAM motif (Fig. 3e). Our findings suggest that better CRISPR sgRNA design should avoid extreme G content in the non-seed region.

**Figure 4.**
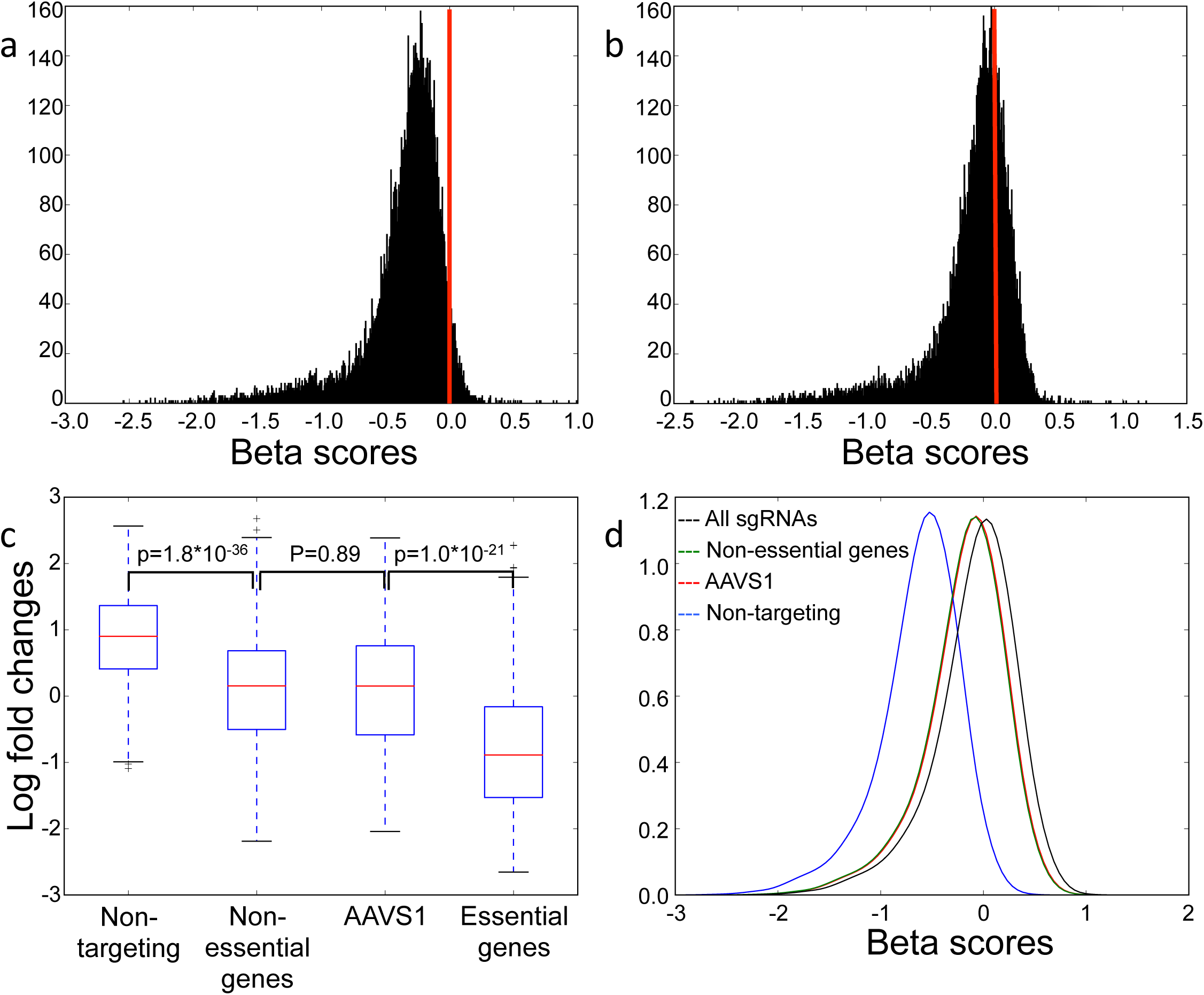
Normalizing read counts using sgRNAs targeting non-essential genes or AAVS1. (a-b) The distribution of beta scores using non-targeting sgRNAs (a) and sgRNAs targeting non-essential genes (b) for normalization. (c) The log fold change distribution of 349 non-targeting sgRNAs, 467 non-essential genes-targeting sgRNAs, 133 AAVS1-targeting sgRNAs, and 725 essential genes-targeting sgRNAs. P values were calculated using two-sided Student’s t-test. (d) The distribution of beta score using all sgRNAs (black), non-essential genes-targeting sgRNAs (green), AAVS1-targeting sgRNAs (red), and non-targeting sgRNAs (blue) for read counts normalizations, respectively.

### SgRNAs targeting AAVS1 or non-essential genes as negative controls reduce false positives in the screen

Correct interpretations of genome-wide screens require proper read count normalization. Since most sgRNAs should generate knockouts without causing phenotype, a straightforward approach is to normalize based on the total read counts of all sgRNAs^35^ (‘total normalization’). Alternatively, many screen libraries include ‘non-targeting’ negative control sgRNAs, which match nowhere in the genome, for normalization (‘non-targeting sgRNA normalization’). In public datasets^5, 11^, ‘total normalization’ resulted in a beta-score distribution centered on zero (Supplementary Fig 6), while ‘non-targeting sgRNA normalization’ led to a skewed distribution of beta scores and most of the genes seemed to be negatively selected (Fig 4a). The bias of ‘non-targeting sgRNA normalization’ is introduced when sgRNAs targeting non-essential genes still impede cell growth from genome cleavage toxicity^29, 36^, regardless of the gene knockout effects. Therefore, a more appropriate choice of negative controls should be sgRNAs that make cleavages at non-essential DNA regions. Indeed, when normalizing read counts using sgRNAs targeting the ‘gold standard’ 927 non-essential genes previously derived from pooled shRNA screens^26^, the beta score distribution is centered on zero (Fig. 4b).

In genome-wide screens, normalizations using either sgRNAs targeting non-essential genes or all genes lead to similar results (Fig. 4b, Supplementary Fig. 6), as the majority of the genes are assumed to be non-essential. Such assumption may fail in focused (or custom) screens where many targeted genes may be under selection, which necessitates the selection of better negative control sgRNAs. AAVS1 (adeno-associated virus integration site 1) has long been recognized as a “safe harbor” site preferred for gene knockins^37, 38^. This region appears to be epigenetically open for efficient cleavage, yet cutting or modification at this site results in no phenotypic changes^39^. To test whether sgRNAs targeting AAVS1 could serve as good negative controls, we first designed a genome-wide screen library containing 134 AAVS1-targeting sgRNAs, 349 non-targeting sgRNAs, as well as 5 sgRNAs per gene in the human genome, and performed screening in a prostate cancer LNCaP-abl cell line. SgRNAs targeting AAVS1 or non-essential genes induced similar LFCs that are stronger than non-targeting sgRNAs, confirming the existence of cleavage toxicity in non-essential regions (Fig. 4c). Also, by comparing normalization methods using different sets of sgRNAs (all, non-targeting, AAVS1-targeting, and non-essential-gene-targeting sgRNAs, respectively), we found normalization using the AAVS1- and non-essential-genes targeting sgRNAs result in almost identical distribution of beta scores (Fig. 5d). Moreover, both ‘total normalization’ and ‘non-targeting sgRNA normalization’ lead to biases, though to different degrees (Fig. 4d).

**Figure 5.**
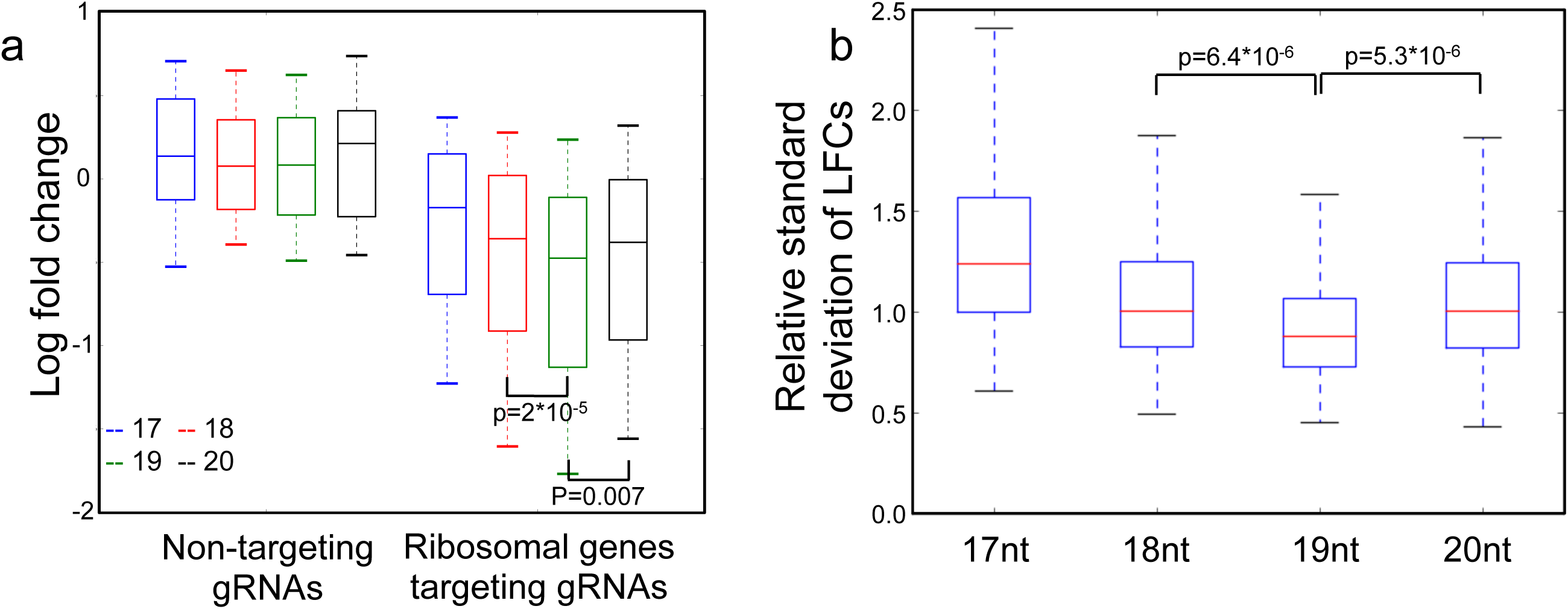
Comparing cleavage efficiencies and signal-to-noise ratios between different lengths of sgRNA spacers. (a) The log fold changes of sgRNAs with spacer lengths ranging from 17- to 20-nts, including non-targeting sgRNAs and sgRNAs targeting ribosomal genes. For each spacer length, there are 100 non-targeting sgRNAs and 1020 ribosomal genes-targeting sgRNAs. P values were calculated using two-sided Student’s t-test. (b) The relative standard deviation of log fold changes of sgRNAs targeting ribosomal genes with spacer lengths ranging from 17- to 20-nts. There are 612 data points (51 ribosomes genes repeated in 12 screens) for each spacer length. P values were calculated using two-sided Student’s t-test.

To evaluate the normalization methods in a focused screen, we also designed a small screening library that targets ~600 genes, including ribosomal genes and well known cancer-related genes (see Methods, Supplementary Tables 2, and 3). The library also includes the same set of AAVS1-targeting and non-targeting sgRNAs. Similar to genome-wide screens, AAVS1-targeting sgRNAs induced stronger negative selections compared with non-targeting sgRNAs (Supplementary figure 7a). Furthermore, using AAVS1-targeting sgRNAs as negative controls in our MAGeCK algorithm greatly increases the sensitivity of the screen, while keeping the same level of false positives (Supplementary figure 7b). These results validated the applicability of including AAVS1-targeting sgRNAs in genome-wide, and more importantly in focused screen libraries.

### 19nt spacers give rise to higher cutting efficiencies and better signal-to-noise ratio

In spCas9 gene editing systems, truncated sgRNAs have been reported to have a better cleavage specificity compared with full-length sgRNAs^40^. However, the performance of truncated sgRNAs in screens compared with full-length sgRNAs remains un-determined. Therefore, in our small screening library, we also designed spacer lengths ranging from 17- to 20-nts for each ribosomal gene and AAVS1-targeting sgRNAs (see Methods). We found that 19nt sgRNAs give rise to significantly stronger LFCs in ribosomal genes, reflecting higher cleavage efficiencies (Fig. 5a). If we use the difference between positive-control sgRNAs (sgRNAs targeting ribosomal genes) and negative-control sgRNA (AAVS1-targeting sgRNAs) as a metric for signal-to-noise, 19nt spacers gave better performance (Supplementary Fig. 8) in 11 of 12 screens. Moreover, for each ribosomal gene, 19nt sgRNAs gave lower relative standard deviation (*i.e.* standard deviation divided by mean) of LFCs, indicating a more stable behavior (and potentially less off-target cleavages) of gene knockout effects (Fig. 5b).

### A new genome-wide library Improved screen performance

Using the rules we uncovered in this study and our previous work^15^, we designed two sub-libraries that target 18,493 human coding genes (named “H1” and “H2”; Supplementary Tables 4 and 5). Each sub-library includes sgRNAs with 19nt-long spacers, and contains 134 AAVS1-targeting sgRNAs, 349 non-targeting sgRNAs, as well as 5 sgRNAs targeting each gene in the human genome. After removing sgRNAs that are enriched in G-nucleotide (>40%) and have perfect matches to other coding regions, we prioritized the remaining sgRNAs based on their predicted cleavage efficiencies^15^ and the number of perfect matches in the whole genome (see Methods). We conducted screens in LNCaP-abl and T47D cell lines using the H1/H2 library and another popular genome-wide library, GeCKO2^4^. We found that the pan-essential genes^26^ are more negatively selected in either H1 or H2 libraries (5 sgRNAs per gene in each library) in both cell lines compared to GeCKO2 (6 sgRNAs per gene) (Supplementary Fig. 9), indicating an improved library performance using our new design rules.

## Discussion

CRISPR-Cas9 knockout screen has been used to systemically interrogate the functions of coding genes and non-coding elements, but data analysis and library design are still in their early stage. We presented MAGeCK-NEST, a computational algorithm to improve the power of CRSIPR screens by incorporating protein-protein interaction (PPI) network into the gene calling procedure. MAGeCK-NEST also employs a maximum-likelihood approach to identify and remove outlier gRNAs, and provides QC metrics to evaluate the quality of the screens. MAGeCK-NEST not only improved CRISPR screen analysis accuracy, but also revealed some important factors in designing better CRISPR-Cas9 screen libraries.

First, we applied MAGeCK-NEST to public genome-wide screen data and identified a set of sgRNA outliers and their sequence characteristics: a higher G-nucleotide counts especially in regions distal from PAM motif. Unexpectedly, the effect of the outliers is independent on the count of C-nucleotide, different from previously studies that suggest the role of ‘GC’ content in determining cleavage efficiencies^5, 17^. Since G-C hybridization strengths in DNA-RNA and RNA-DNA hybrids are similar, the distinct effect of G- and C-nucleotides suggests a more crucial role of DNA-endonuclease rather than DNA-RNA interaction in determining off-target effects. Moreover, these sgRNAs do not match to other genomic regions^11^, and the potential off-target cleavages induced by these sgRNAs may occur less frequently at regions not predictable by sequence similarity, and regions less likely to be detected using current off-target detection technologies^41, 42^. Second, we found normalization using non-targeting sgRNAs, as compared to using all sgRNAs or sgRNAs targeting non-essential genes, leads to higher false positive rates. This might be because cleavages in non-essential regions can still induce toxicity in cell growth, in consistence with two recent studies showing false positive hits from highly amplified regions in cancer genomes^29, 36^. Through CRISPR screening experiments, we confirmed that sgRNAs targeting AAVS1 or non-essential genes could serve as better negative controls and result in fewer false positives compared with non-targeting sgRNAs. Finally, we discovered that 19nt sgRNAs consistently provide better cleavage efficiencies and signal-to-noise separations compared with other lengths (17, 18, 20nt). Therefore, using 19nt sgRNAs in either low-throughput experiments or high-throughput screens may gives rise to a more accurate inference of gene knockout effects.

Although we characterized multiple features of CRISPR screens using computational approaches, the exact mechanisms behind these findings remain unknown. First, it is unclear how sgRNAs with higher G-nucleotide content are associated with stronger outliers. We suspected that outlier gRNAs with high G-nucleotides have promiscuous off-target binding and cutting at many CpG islands in the genome. Existing experimental approaches to detect off-target cleavages^41, 42^ may be limited to study these gRNAs, as the cleavages in each binding site may be low. Second, although we have shown the advantages of using 19bp sgRNA spacers from statistical perspectives, how different lengths of sgRNA spacers give rise to various cleavage strengths and off-targets remain to be determined. Last but not least, all the above findings are derived in SpCas9 system, and the rules in different RNA-guided DNA endonuclease systems require further investigations.

Collectively, our study provided novel insights into both the analysis and design of CRISPR screens. We designed two genome-wide libraries using the rules we uncovered, and demonstrated their better performances compared to GeCKO2. The algorithm, the design rules, as well as the libraries, will benefit and expedite application of CRISPR techniques.

## Method

### Formulating predicted beta scores using PPI

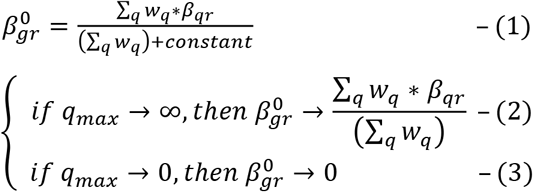

The predicted beta score of gene *g* in condition *γ*, 
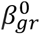
, was derived by weighted average of the beta scores of interacting gene *q* in condition *γ*, *β*_*qr*_, with weighting to represent interacting strength, *w_q_*, provided by the String v9.1^43^. However, such formulation of 
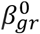
 should be modified to meet certain desired characteristics. First, when there are only a few interacting genes, the derived predicted beta scores becomes unreliable. Therefore, the 
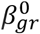
 should approximate to weighted average of *β*_*qr*_ when number of interacting genes increases, but gets closer to zero when number of interacting genes decreases (Equation (2–3)). This requirement is fulfilled by adding a positive *constant* in Equation (1). Second, in order to use 
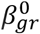
 as a prior in estimating *β*_*gr*_, an ideal 
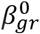
 should be an unbiased estimator of *β*_*gr*_. To fulfill these two requirements, we used published screen data and determined that when the number of sgRNAs per gene is between 4 and 10, using 3 as *constant* would allow the predicted beta scores become unbiased estimator of observed beta scores.

### Adopting predicted beta score as Bayesian prior to estimate beta score

Similar to MAGeCK-VISPR^24^, the read count of sgRNA *i* in sample *j*, or *K_ij_*, is modeled as:

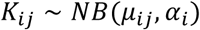

Where *μ_ij_ and α_i_* are the mean and over-dispersion factor of the negative binomial (NB) distribution, respectively. The mean value *μ_ij_* is further modeled as:

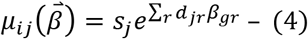

Where *s_j_* is the size factor of sample *j* for adjusting sequencing depths of the samples. To deal with complex experimental settings, we included design matrix (*D*). With *J* samples affected by *R* conditions, *D* is a binary matrix with its element *d_jr_* = 1 if sample *j* is affected by condition *r*, and 0 otherwise. In order to incorporate 
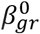
 in Bayesian framework to estimate *β*_*gr*_, we reformulated the goal function to a regularization form:

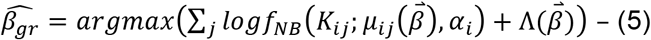

Where

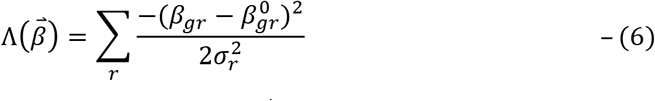

In Equation (5), the regularization term, 
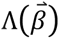
, draws 
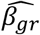
 closer to the prior mean, 
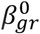
, and the amount of movement depends on the *observed Fisher information* provided by the sgRNAs. In Equation (6), we assumed the empirical prior of *β*_*qr*_ follows a normal distribution centered at 
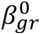
.

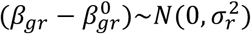

The width of the prior distribution, *σ_r_*, was calculated using the naive estimators of *β*_*rr*_. For robust estimator of *σ_r_*, we adopted quantile matching: the standard deviation *σ_r_* is chosen such that (1-p) empirical quantile of the absolute value of the observed beta scores matches the (1-p/2) theoretical quantile of normal distribution *N*(0, *σ*^2^), and set default p value as 0.05:

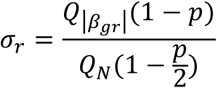

To solve Equation (5), we re-formulated the equation as the function of 
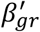
. Assume:

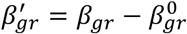

Then Equation (4) can then be re-written as:

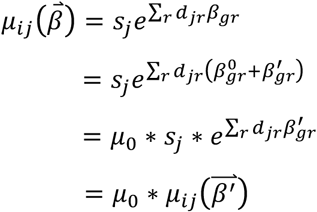

Where *μ*_0_ is a constant:

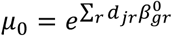

Then Equation (5) can thus be re-written as:

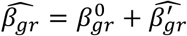

Where

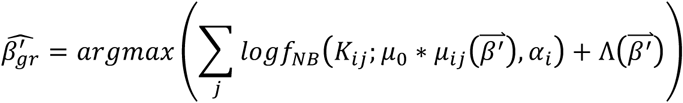

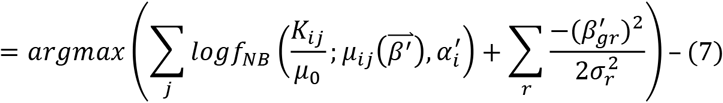

In order to make

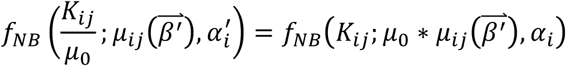

The transformed over-dispersion factor, 
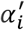
, can be deducted as:

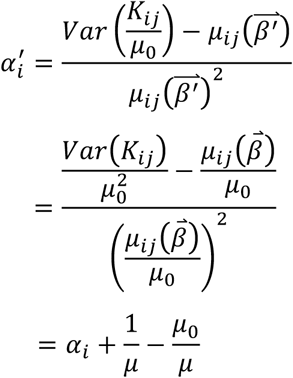

Now the re-formulated Equation (7) can be solved using the iteratively reweighted ridge regression algorithm as described previously^24, 34, 35^.

### Estimating over-dispersion factors

The calculation of sgRNA-wise over-dispersion factor, *α*, is similar to what DESeq2 adopted^35^. Specifically, the over-dispersion factor of sgRNA *i*, *α_i_*, was obtained via maximizing the Cox-Reid adjusted likelihood of the dispersion.

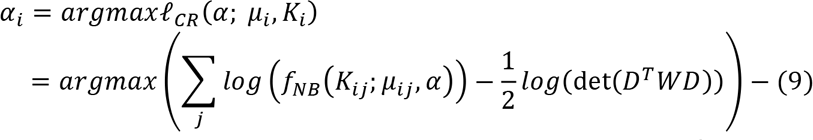

The second term provides Cox-Reid bias adjustment, where *W* is the diagonal matrix with its values given by 
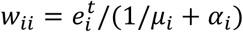
. The equation (9) could then be solved using stepwise descent along *logα* as previously described^35^.

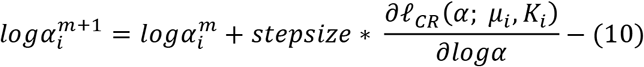

The derived sgRNA-wise over-dispersion factors were then used to fit the trend function:

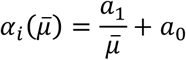

### Identifying sgRNA outliers

Single outlier that does not fit the assumed distributions can overly influence the estimations of beta score. Therefore, we tried to identify these outliers using 3-step approach: candidate outlier prediction, candidate outlier validation, and outlier detection.

#### Step-1: Candidate outlier prediction

An sgRNA is likely to be an outlier if its log fold is extremely different from other sgRNAs. Therefore, in the first step, candidate outlier prediction, we identified the potential sgRNAs outliers by considering their log fold changes (LFCs). For each paired conditions, we calculated the median and standard deviation of the LFCs, and defined the candidate outliers if their LFCs fall beyond median ± 1.5 standard deviation. To make the standard deviation estimator robust against extremely high absolute LFCs, we used quantile matching with p set by default to 0.34.

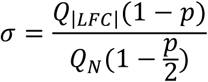

#### Step-2: Candidate outlier validation

Noticing that an sgRNA outlier may significantly influence the beta score estimation, a candidate outlier is validated if there is a significant change of *β*_*ir*_ after removing the candidate outlier. Therefore, in the second step, the candidate outlier validation, we calculated the beta score with and without the candidate outlier respectively using Equation (5). Define:

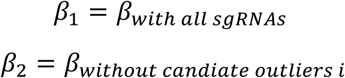

Then candidate outlier *i* is validated if:

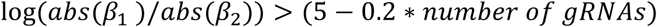

With outlier removal, we could prevent the beta score estimation from distortion by strong outliers.

#### Step-3: Outlier detection

With previous 2 steps, we could estimate the beta scores robustly. However, some moderate outliers remain un-identified if sufficient sgRNAs prevent the beta score from distortion by single outlier. Therefore, with robustly estimators of beta scores, in the final step we re-defined an sgRNA as an outlier if the likelihood of observing its count and corresponding beta score falls below certain threshold. The threshold was determined using two strategies. The first one is using the validated outliers as “flags”, in which the threshold is determined so that 90% of validated outliers defined in step 2 can be removed. In the second strategy, we directly assigned an sgRNA as outlier if its likelihood is in the lower 5% of all sgRNAs, a percentage same as what we observed in recently published screens (Fig. 3a).

### Extracting sequence features using Elastic-Net regression

To identify the sequence features that are associated with stronger sgRNA outliers, we applied Elastic-Net regression to extract the sequence features as our previous work^15^. Suppose *X* = {*X*_1_, *X*_2_,…, *X*_*n*_} is the set of encoded sequence vectors and *Y* = {*Y*_1_, *Y*_2_,…, *Y*_*n*_} is the set of outputs representing whether the sgRNAs are stronger outliers, where *n* is the number of sgRNAs samples for training. Let *M* be the length of the input vectors, the Elastic-Net regression computes the parameters 
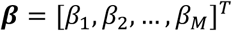
 that minimizes an object function E:

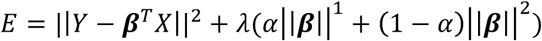

Where *α* and *λ* are parameters estimated using cross validation, 
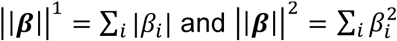
. We used glmnet in R package to implement the Elastic-Net regression^34^.

### Representation of the regression coefficients

We used Seq2Logo 2.0 server (http://www.cbs.dtu.dk/biotools/Seq2Logo/) to illustrate the coefficients derived from Elastic-Net regression^44^.

### D-distance statistic

The Kolmogorov–Smirnov test (K-S test) is a nonparametric test of the equality of continuous, one-dimensional probability distributions that can be used to compare two samples. We used the D-distance statistic in K–S test to measure how different spacer lengths affect the separation between ribosomal genes-targeting sgRNAs (positive controls) and AAVS1-targeted sgRNAs (negative controls).

### Customized-design libraries

We designed a small library to compare the normalization using AAVS1-targeting and non-targeting sgRNAs, as well as how spacers with different lengths ranging from 17nts to 20nts influence the cleavage efficiencies (Supplementary Table 2). The library contains four major categories of sgRNAs: AAVS1-targeting sgRNAs, non-targeting sgRNAs, sgRNAs targeting 51 ribosomal genes and 503 cancer-related genes, which were selected using published cancer signatures^45–47^. Detailed designs are summarized as following table.

| Target | Spacer lengths | Number of each gene-length | Total number of sgRNAs |
| --- | --- | --- | --- |
| AAVS1 | 17-20 | 204 | 1632 |
| Non-targeting | 17-20 | 100 | 400 |
| 51 ribosomal genes | 17- 20 | 20 | 4080 |
| 503 selected cancer-related genes | 20 | 12 | 6036 |

The oligos were synthesized at CustomArray©.

### Design of H1 and H2 libraries

We designed two genome-wide screen libraries, H1 and H2, using the rules we uncovered. Totally 185,634 19-nt sgRNAs were designed, targeting 18,453 genes in human, as well as AAVS1-targeting and non-targeting sgRNAs. We identified all candidate sgRNAs by locating NGG protospacer adjacent motifs (PAMs) in coding regions on both strands. In order to maximize the on-target cleavage efficiency, we scored each candidate sgRNA using sequence-based cleavage efficiency predictive model^15^. To mitigate the potential off-targets (or the resulting outliers), we adopted 19-nt sgRNAs, removed the common SNP/mutant loci, counted the G-nucleotide counts, and estimated the potential off-targets by considering the sequence uniqueness. Detailed steps are as follows:

Filter stage:

1. Select all 19bp sequences upstream of the “NGG” PAM motif, in the coding regions of the target gene;
2. Remove the sequences that:
  i. hit SNP / mutant loci;
  ii. with > 40% of G;
  iii. with off-target perfect match in the genome;
3. Rank the remaining sequences in descending order of predicted efficiency score.
4. If the number of remaining sequences is smaller than 10, go to Rescue stage, otherwise select the top 10 sequences with highest predicted efficiency scores to be sgRNA targets.

Rescue stage:

1. Select all remaining sequences in the Filter stage to be sgRNA targets;
2. Rescue the sequences with off-target perfect match in non-coding regions but not in coding regions;
3. Rank the sequences rescued in 2) in ascending order of the number of off-target perfect matches. If two or more sequences has the same number of off-target matches, rank them in descending order of efficiency score;
4. Add the rescued sequences in 2) to the sgRNA target list in order of the ranks in 3) until the list has a size of 10, or all the rescued sequences are added. If the target list has a size of 10, exit the Rescue stage.
5. Rescue the sequences with off-target perfect match in coding regions;
6. Rank the sequences rescued in 5) in ascending order of the number of off-target matches in the genome. If two or more sequences has the same number of off-target matches, rank them in descending order of efficiency score;
7. Add the rescued sequences in 6) to the sgRNA target list in order of the ranks in 6) until the list has a size of 10, or all the rescued sequences are added.

Finally, we separated all sgRNAs evenly to H1 and H2 sub-libraries as following table:

|  | Gene-targeting sgRNAs | Non-targeting sgRNAs | AAVS1-targeting sgRNAs | Total sgRNRs |
| --- | --- | --- | --- | --- |
| H1 | 92,287 | 398 | 133 | 92,817 |
| H2 | 92,285 | 399 | 134 | 92,817 |

The oligos were synthesized at CustomArray©.

### Cell lines and cell culture

LNCaP-abl (abl) cell line was provided by Zoran Culig (Innsbruck Medical University, Austria). The abl cells were cultured in the RPMI 1640 phenol red-free medium supplemented with 10% charcoal/dextran-treated fetal bovine serum (FBS), 2mM glutamine, 100 ug/ml penicillin and 100units/ml streptomycin for the experiments. The T47D cells obtained from the American Type Culture Collection were maintained in RPMI 1640 phenol red medium plus 10% FBS. The 293FT cells bought from ThermoFisher were cultured in DMEM media supplemented with 10% fetal bovine serum, glutamine and penicillin-streptomycin.

### Plasmid construction and lentivirus production

The sgRNA library synthesized at CustomArray© were amplified by PCR as previously described (PMC4089965). The PCR products were subsequently ligated into lentiCRISPR V2 plasmid, followed by transformation to competent cells for amplification according to a online protocol (GeCKO library Amplification Protocol from Addgene). After library plasmid had been amplified, we isolated the plasmid and construct a sequencing library for Miseq to ensure library diversity. To make lentivirus, T-225 flasks of 293FT cells were cultured at 40%~50% confluence the day before transfection. Transfection was performed using X-tremeGENE HP DNA Transfection Reagent (Roche). For each flask, 20 ug of lentivectors, 5 ug of pMD2.G, and 15 ug of psPAX2 (Addgene) were added into 3 ml OptiMEM (Life Technologies). 100 ul of X-tremeGENE HP DNA Transfection Reagent was diluted in 3 ml OptiMEM and, after 10 min, it was added to the plasmid mixture. The complete mixture was incubated for 20 min before being added to cells. After 6 h, the media was changed to 30 ml DMEM + 10% FBS. After 60 h, the media was removed and centrifuged at 3,000 rpm at 4 °C for 10 min to pellet cell debris. The supernatant was filtered through a 0.45 um low protein binding membrane. The virus was ultracentrifuged at 24,000 rpm for 2 h at 4 °C and then resuspended overnight at 4°C in DMEM + 10% FBS. Aliquots were stored at –80°C.

### CRISPR screens

Cells of interest were infected at a low MOI (0.3～0.5) to ensure that most cells receive only 1 viral construct with high probability. To find optimal virus volumes for achieving an MOI of 0.3–0.5, each new cell type and new virus lots were tested by spinfecting 3×106 cells with several different volumes of virus. Briefly, 3×106 cells per well were plated into a 12 well plate in the appropriate standard media for the cell type (see below) supplemented with 8 ug/ml polybrene. For T47D cells, standard media is RPMI 1640 supplemented with 10 % FBS. Each well received a different titrated virus amount (usually between 5 and 50 ul) along with a no-transduction control. The 12-well plate was centrifuged at 2,000 rpm for 2 h at 37°C. After the spin, media was aspirated and fresh media (without polybrene) is added. Cells are incubated overnight and then enzymatically detached using trypsin. Cells are counted and each well is split into duplicate wells. One replicate receives 4ug/mL puromycin for abl cells or 3.5 ug/ml puromycin for T47D cells. After 3 days (or as soon as no surviving cells remained in the no-transduction control under puromycin selection), cells were counted to calculate a percent transduction. Percent transduction was calculated as cell count from the replicate with puromycin divided by cell count from the replicate without puromycin multiplied by 100. The virus volume yielding a MOI closest to 0.4 was chosen for large-scale screening.

For the screens using ~600 gene library, spin-infection of 2×107 cells were performed by one 12-well plates. And large-scale spin-infection of 2×108 cells was carried out using four of 12-well plates with 4×106 cells per well for a genome-wide screen. Wells are pooled together into larger flasks on the day after spinfection. After three days of puromycin selection, the surviving cells (T47D and abl) were divided into two groups: one group for 0 day control, and the other one was cultured in RPMI or DMEM medium plus 10% FBS for four weeks before genomic DNA extraction and analysis. Two rounds of PCR were performed after gDNA had been extracted, and 300ug DNA per sample was used for library construction. Each library was sequenced at 3~30 million reads to achieve ~300X average coverage over the two different CRISPR libraries. The 0 day sample library of each screen could serve as controls to identify positively or negatively selected genes or pathways.

### PCR primers for library construction

The first round of PCR:

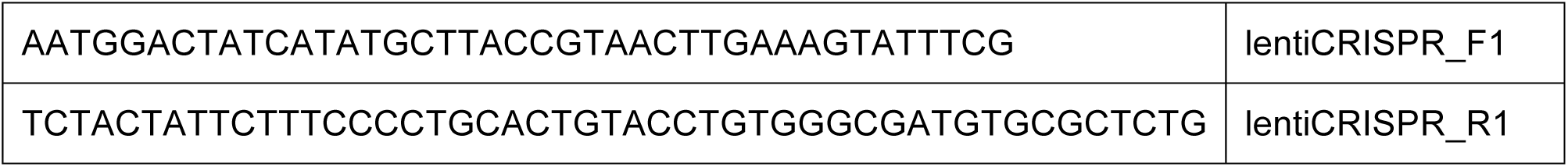

The second round of PCR:

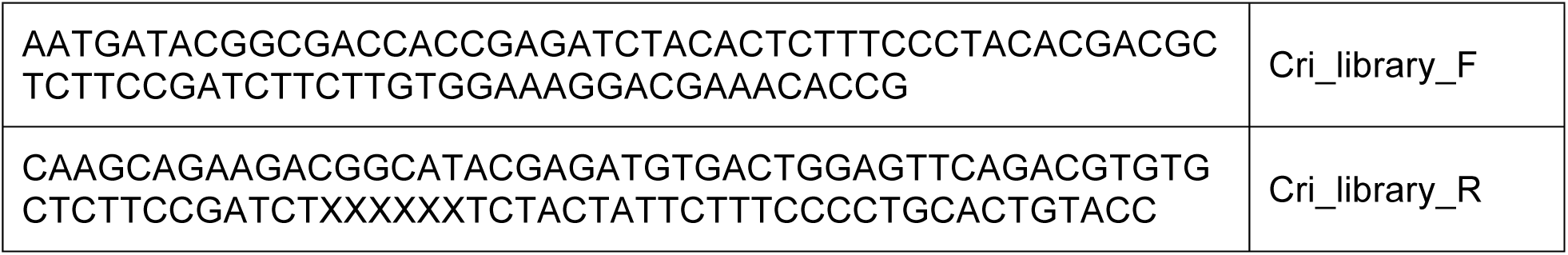

(XXXXXX denotes the sample barcode)

Sequencing primer (read1): GCTCTTCCGATCTTCTTGTGGAAAGGACGAAACACCG

Indexing primer: CATCGCCCACAGGTACAGTGCAGGGGAAAGAATAGTAGA

### Code availability

The MAGeCK-NEST workflow is available open source at https://bitbucket.org/liulab/mageck_nest under the MIT license.

### Author contributions

X.S.L. and M.B. conceived and supervised the whole project. C.H.C, W.L. developed the algorithm, C.H.C, W.L., T.X., and H.X. designed the screen and performed the analysis. T.X performed the experiments. P.J., C.M., and M.B. helped with technical clarifications and result interpretations. C.H.C, W.L., T.X. and X.S.L. wrote the manuscript with help from all other authors.

